# Brain Iron as a Surrogate Biomarker of Pathological TDP-43 Identifies Brain Region-Specific Signatures in Ageing, Alzheimer’s Disease and Amyotrophic Lateral Sclerosis

**DOI:** 10.1101/2025.10.02.680028

**Authors:** Fergal M. Waldron, Holly Spence, Orjona Stella Taso, Fiona L. Read, Irika R. Sinha, Katherine E. Irwin, Philip C. Wong, Jonathan P. Ling, Jenna M. Gregory

## Abstract

**Background:** TDP-43 pathology is a defining feature of several neurodegenerative diseases, but its prevalence and regional distribution in ageing and disease are not well characterised. We investigated the burden of brain TDP-43 pathology across ageing, Alzheimer’s disease (AD), and amyotrophic lateral sclerosis (ALS), and examined ferritin as a region-specific correlate of TDP-43 pathology.

**Methods:** Pathological TDP-43 was detected using an *HDGFL2* cryptic exon *in situ* hybridisation probe and a TDP-43 RNA aptamer, providing greater sensitivity and specificity than antibody-based approaches. Amygdala, hippocampus, and frontal cortex tissue was analysed from non-neurological controls (ages 40–80), AD cases, and ALS cases. Ferritin (as a proxy for iron accumulation) was quantified in parallel to assess its association with TDP-43 pathology.

**Findings:** TDP-43 pathology was detectable from the fourth decade of life, with a 4.5-fold increase in hippocampal involvement after age 60 years. In AD, pathology was present in 90% of cases and distinguished from ageing by selective amygdala involvement. In ALS, TDP-43 pathology was nearly ubiquitous across all regions studied. Regional ferritin strongly predicted TDP-43 burden: amygdala ferritin explained 87% of TDP-43 variance in ALS and 66% in AD, while hippocampal ferritin differentiated AD from controls. Across AD, ferritin explained between 43–81% of regional TDP-43 variance.

**Interpretation:** TDP-43 brain pathology emerges in midlife with increased involvement after age 60 years, exhibits disease-specific regional signatures in AD and ALS, and is closely linked to ferritin accumulation. As TDP-43 confers a worse prognosis in AD, the capacity of ferritin, detectable with iron-sensitive MRI, to serve as a proxy for regional TDP-43 burden highlights its promise as a biomarker for disease stratification and prognosis.

**Short Abstract:** Here we show that pathological TDP-43 emerges during normal ageing from the fourth decade of life, with a 4.5-fold increase in hippocampal involvement after 60 years. In Alzheimer’s disease (AD), TDP-43 pathology was present in 90% of cases and distinguished from ageing by disproportionate amygdala involvement, while in amyotrophic lateral sclerosis (ALS) it was nearly ubiquitous across hippocampus, amygdala, and frontal cortex. Using sensitive detection tools, we demonstrate that region-specific ferritin strongly predicts TDP-43 burden: amygdala ferritin explained 87% of variance in ALS and 66% in AD, while hippocampal ferritin differentiated AD from controls. Across AD, ferritin levels in all three regions explained 43–81% of TDP-43 variance. As TDP-43 pathology confers a worse prognosis in AD, the ability of ferritin, quantifiable with iron-sensitive MRI, to serve as a proxy for regional TDP-43 burden highlights its potential as a biomarker for disease stratification and prognostic assessment.

**Graphical Abstract:** 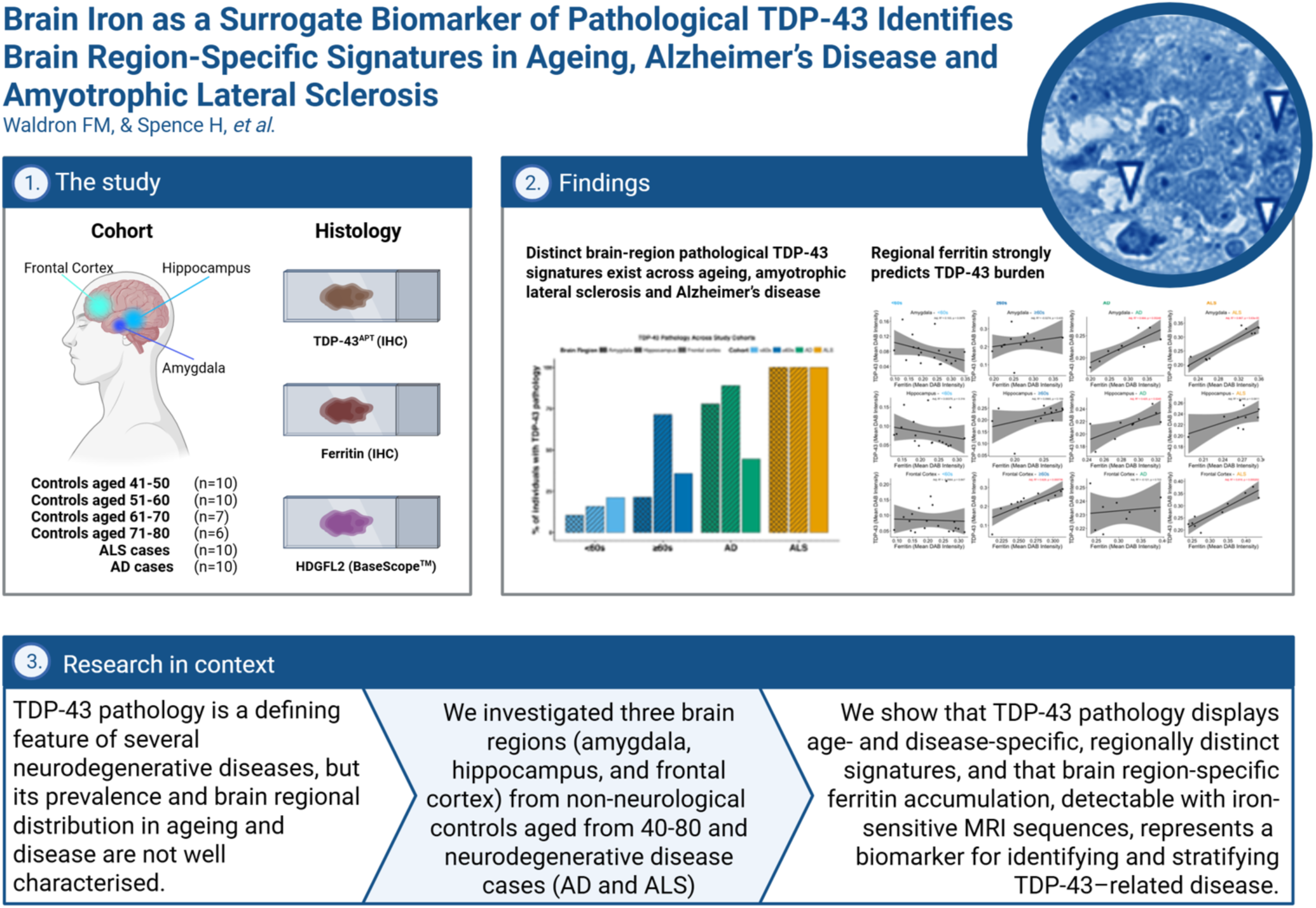

**Highlights:**

- TDP-43 brain pathology occurs in normal ageing from early in the fourth decade, characterised by a 4.5-fold increase in hippocampus pathology from the sixth decade.
- TDP-43 brain pathology is detectable in 90% of AD cases, with a disease-signature of increased amygdala pathology relative to age-matched controls.
- In ALS, TDP-43 is nearly ubiquitous in amygdala, hippocampus and frontal cortex.
- Hippocampus high brain ferritin distinguishes AD from and age-matched controls
- Brain ferritin is a brain region-specific marker of TDP-43 pathology in ageing and disease, with amygdala ferritin explaining 87% of the variance in amygdala TDP-43 pathology in ALS, and 66% of amygdala TDP-43 pathology in AD
- In AD, ferritin levels for all three brain regions explain between 43-81% of variance in their TDP-43 pathology levels

## Introduction

Transactive response DNA-binding protein 43 (TDP-43) has emerged as an important pathological hallmark in a wide spectrum of neurodegenerative diseases, most notably amyotrophic lateral sclerosis (ALS) and frontotemporal lobar degeneration (FTLD). Beyond these conditions, growing evidence suggests that TDP-43 pathology is also present in other disorders, including Alzheimer’s disease, hippocampal sclerosis of ageing, and chronic traumatic encephalopathy. Despite this broad involvement, the extent and clinical significance of TDP-43 pathology are likely underestimated. A role for TDP-43 in normal ageing has also been recognized, with pathology detected in a subset of cognitively normal older individuals ^1,2^. This observation has given rise to the concept of limbic-predominant age-related TDP-43 encephalopathy (LATE), highlighting that TDP-43 contributes not only to classical neurodegenerative syndromes but also to age-related cognitive decline. ^3,4^. However, the distribution, burden, and impact of TDP-43 pathology across ageing and disease remains incompletely characterised. Importantly, there is a growing need to appreciate the extent of TDP-43 pathology beyond the context of traditional neurodegenerative disease silos to determine if moving from clinical syndromes to biological definitions of disease (as with Alzheimer’s disease ^5^, Huntington’s disease ^6^, and neuronal α-synuclein disease ^7^) could be appropriate for TDP-43 proteinopathies ^8^.

One factor limiting progress is methodological. Many prior studies have relied on conventional antibody-based detection methods, which lack sufficient sensitivity and may therefore underestimate the true burden of pathology. This limitation obscures both the prevalence of TDP-43 pathology and its association with clinical outcomes. In addition, the contribution of TDP-43 in the context of co-existing proteinopathies remains poorly understood, despite its importance for clarifying disease mechanisms. For instance, in Alzheimer’s disease (AD), TDP-43 pathology is linked to increased brain pTau burden and worse clinical prognosis. These insights highlight that the implications of TDP-43 extend beyond prognostication, with therapeutic relevance not only for ALS but also for other TDP-43 related neurodegenerative diseases such as LATE ^9^, AD ^10,11^, and AD with hippocampal sclerosis ^12^.

There is a pressing need for functional and accessible biomarkers of TDP-43 pathology that can operate across blood^13^, CSF, tissue, and imaging platforms. Evidence of TDP-43 loss of function, such as cryptic exon incorporation observed in Alzheimer’s disease (AD) brains that lack visible TDP-43 inclusions but show nuclear clearance of TDP-43, suggests that current detection tools may lack sufficient sensitivity to detect all relevant aggregation events^14^. As a result, important aspects of TDP-43 pathology may go undetected.

To address this, we have developed a TDP-43 RNA aptamer ^15^ with greater sensitivity and specificity than traditional antibody approaches, optimized for use in the staining of human tissue ^16,17^. Using this TDP-43 RNA aptamer we have shown evidence of novel TDP-43 pathologies such as early nuclear aggregation ^16^, and improved detection of pre-symptomatic, non-central nervous system TDP-43 pathologies in several peripheral tissues ^18^, compared to antibody approaches in the same patient cohort ^19^.

Here, we employ more sensitive tools (RNA aptamer and ISH probes to detect cryptic splicing events) to provide a detailed characterization of TDP-43 brain pathology across the spectrum of normal ageing and in the contexts of AD and ALS. Using post-mortem brain tissue, we focus on regions known to be particularly vulnerable to TDP-43 pathology, the amygdala, hippocampus, and frontal cortex. By systematically comparing these regions across ageing and disease states, we aim to refine estimates of pathological burden, delineate region-specific patterns of involvement, and assess how these patterns differ between neurodegenerative disease and age-related processes. This approach enables a more accurate understanding of the distribution and significance of TDP-43 pathology, with the ultimate goal of clarifying its role in clinical symptoms and disease progression.

Finally, given the lack of reliable *in vivo* markers of TDP-43 pathology, identifying proxy measures that can be captured with existing imaging modalities is of critical importance. Ferritin, the major intracellular iron storage protein, represents a proxy for iron accumulation which can be quantified non-invasively using iron-sensitive MRI techniques and reflects brain region-specific iron accumulation – indeed, such techniques have recently revealed that brain region-specific iron is a strong predictor of mild cognitive impairment and cognitive decline in cognitively unimpaired older adults, especially those with amyloid pathology ^20^. We have shown previously ^21,22^ that iron accumulation (in the form of ferritin) is associated with TDP-43 burden in symptomatic brain regions in ALS but extending this analysis to additional cases and brain regions and to assess this relationship in ageing and AD is clearly warranted, especially given the association between brain iron accumulation and cognitive dysfunction in ageing cohorts ^23,24^. If ferritin levels track with the regional distribution of pathological TDP-43, this would provide a practical and scalable approach to stratifying individuals at risk, distinguishing disease from normal ageing, and predicting prognosis, particularly where TDP-43 pathology is associated with accelerated cognitive decline and worse clinical outcomes in AD ^25^, and with natural ageing ^26^. Establishing ferritin accumulated iron as an imaging correlate of TDP-43 pathology would therefore represent a crucial step toward bridging post-mortem discoveries with clinically actionable biomarkers in ageing and neurodegenerative disease.

## Methods

### Study cohorts

We assembled and analyzed a post-mortem tissue cohort of 53 individuals, balanced for gender (χ^2^=3.189, df=1, p=0.074; Chi-squared test) with 33 males and 20 females.

Amongst the 53 individuals, 33 were people who died of non-neurological diseases aged between 40 and 73 years old at death, and whose brains were collected as part of the sudden death brain bank. Ten of the 53 individuals were diagnosed with ALS, which was confirmed at post-mortem, with a further 10 individuals diagnosed with AD during life, and confirmed at post mortem.

To compare aspects of TDP-43 pathology across ageing and neurodegenerative diseases we established 4 study cohorts (see **Table 1**):

i. a young *“<60s cohort*” consisting of 19 individuals aged 40-59 years (median=50; IQR=47.5-57) at autopsy;
ii. an older “*≥60s cohort*” consisting of 14 individuals aged 60-74 years (median=69; IQR=63.5-71.8) at autopsy;
iii. an “*ALS cohort*” consisting of 10 individuals aged 61-80 years (median=70.5; IQR=65.5-74.5) at autopsy;
iv. an “*AD cohort*” consisting of 10 individuals aged 69-84 years (median=74; IQR=72-77.2) at autopsy;

**Table 1.**
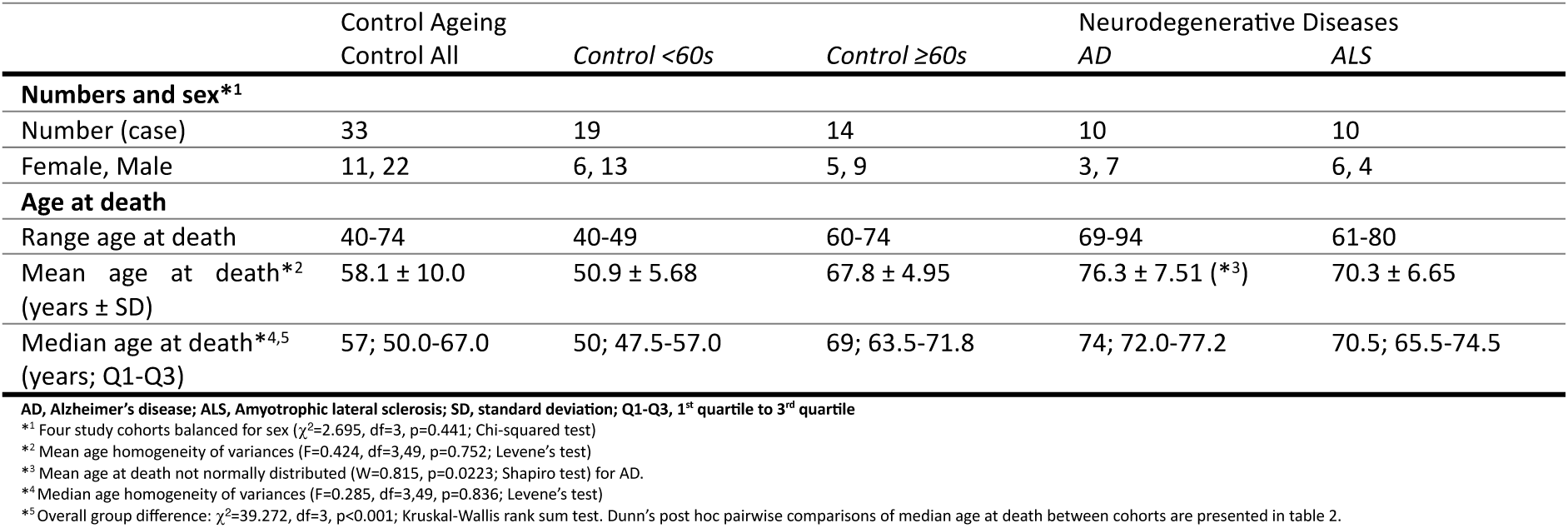
Study cohort demographics. Demographic characteristics of control ageing and neurodegenerative disease cohorts included in the study. Data are presented for all controls, as well as age-stratified control groups (<60s, ≥60s), Alzheimer’s disease (AD), and amyotrophic lateral sclerosis (ALS) cases. Cohort features presented include the number of cases, sex distribution, and age at death.

See figure 1A for an illustration of the study design (with cohorts and histological investigation information) and **Table 1** for a detailed demographic summary.

**Figure 1.**
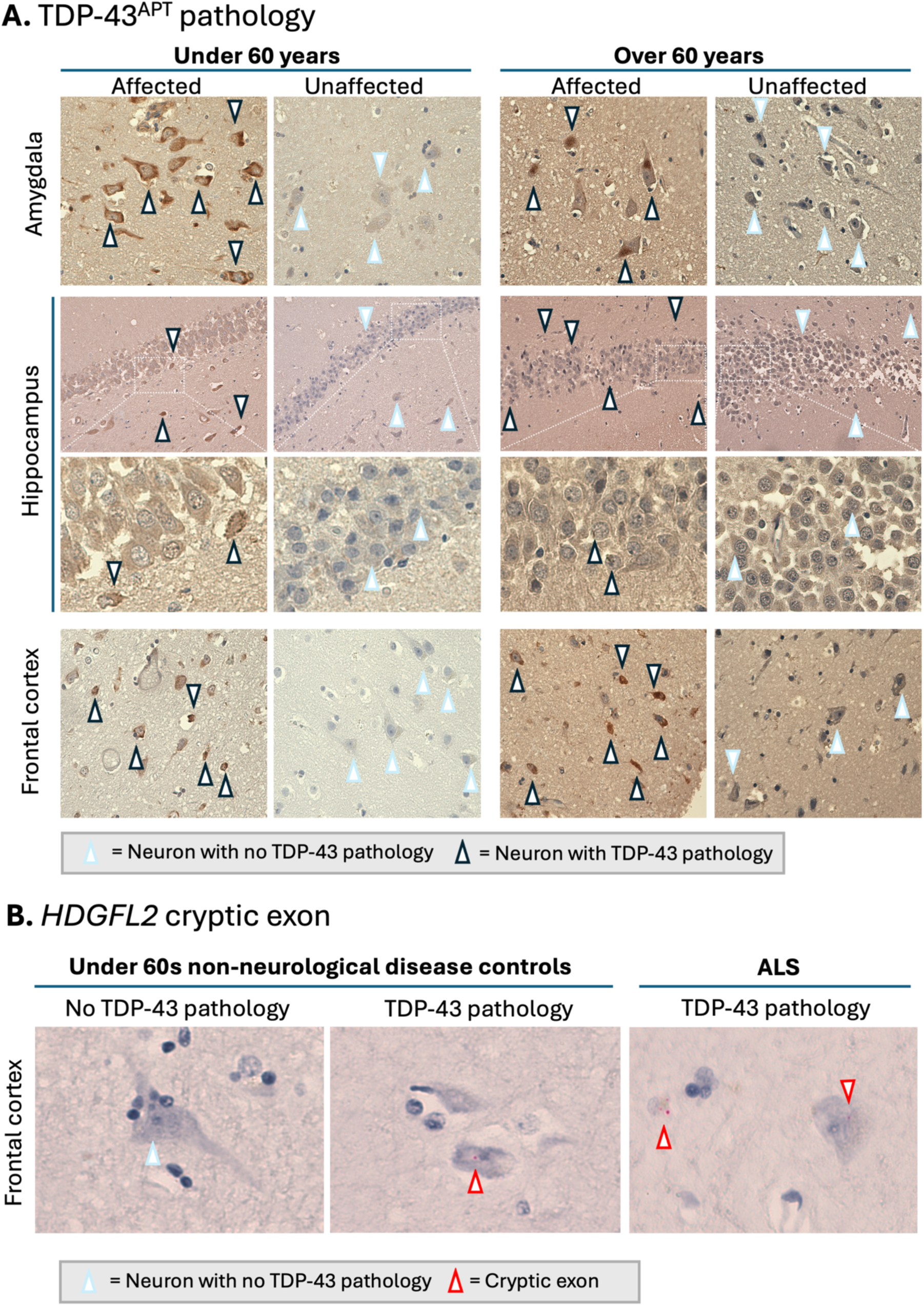
TDP-43 pathology occurs in all 3 brain regions in normal ageing. **A,** Representative images of the amygdala, hippocampus, and frontal cortex from unaffected (controls without TDP-43 pathology) and affected (controls with TDP-43 pathology) individuals, stratified by age (<60 and ≥60 years). TDP-43 aptamer staining highlights neuronal pathology: neurons with no TDP-43 pathology are indicated by arrowheads with a light blue outline, whereas neurons exhibiting TDP-43 pathology are indicated by arrowheads with a black outline. Comparisons across brain regions and age groups reveal region- and age-associated differences in TDP-43 pathology, but with equivalent pathological burden in affected individuals. **B,** Representative images show neurons labelled with an HDGFL2 cryptic exon-specific probe, where each red dot corresponds to a single mRNA transcript containing the cryptic exon, reflecting TDP-43 loss-of-function.

Our *a priori* rationale for dividing our ageing cohorts into *<60s* and *≥60s* was as follows; 1) the median age of at death for control ageing brains was 58 (IQR: 50.0-67.0), so a sixth-decade cut-off facilitated the most statistical power to perform inferential analyses (in addition to investigations of the relationship between normal ageing and TDP-43 pathology) investigating differences in aspects of TDP-43 brain pathology between “younger” and “older” control brains, and 2) There is extensive evidence of significant ageing-related changes in brain health and biology begin to occur in the sixth decade^27–31^.

To understand aspects of TDP-43 pathology across ageing and neurodegenerative diseases, it is necessary to consider the potential confounding factors of sex, and age at death, differences between study groups. The four study cohorts (*<60s, ≥60s, ALS* and *AD*) were balanced for gender (χ^2^=2.695, df=3, p=0.441; Chi-squared test; see **Table 1**). Data exploration of age at death indicated homogeneity of variances for both mean (F=0.424, df=3,49, p=0.752; Levene’s test; see **Table 1**), and median (F=0.285, df=3,49, p=0.836; Levene’s test; see **Table 1**), across all four study cohorts. Amongst the four study groups however, age at death was not normally distributed for AD (W=0.815, n=10, p=0.022; Shapiro test; see **Table 1**). Therefore, medians with interquartile ranges are reported for summaries of age at death for study cohorts, and (conservatively, and for consistency) non-parametric statistics were employed throughout when comparing age distributions between study cohorts.

As expected, median age at death differed between study groups (χ^2^=39.272, df=3, p<0.001; Kruskal-Wallis rank sum test; see **Table 2**; **Suppl. Fig. 1A**). Pairwise Dunn’s post hoc statistics comparing median age at death between cohorts are presented in **Table 2** - there were no significant differences in age at death between *ALS* and *AD* (z=1.051, p=0.880; Dunn’s test), but also between *≥60s* and *ALS* (z=0.678, p=1; Dunn’s test), and *≥60s* and *AD* (z=1.813, p=0.210; Dunn’s test). Age at death was significantly lower for the *<60s* cohort compared with *≥60s* (z=4.017, p<0.001; Dunn’s test), but also for both *ALS* (z=4.340, p<0.001; Dunn’s test) and *AD* (z=5.543, p<0.001; Dunn’s test) (see **Table 2**; **Suppl. Fig. 1A**).

**Table 2.** Pairwise comparisons of median age at death between study cohorts. Median age at death (years; Q1– Q3) is presented for each pairwise comparison, with overall differences in median age at death assessed using the Kruskal–Wallis rank sum test, with Dunn’s post hoc test applied for pairwise comparisons, with p-values adjusted for multiple comparisons. Test statistics (z) and corresponding p-values are reported, with significant differences indicated by asterisk (*).

|  | Test Statistic (z) | p-value |
| --- | --- | --- |
| Control <60s vs. Control ≥60s<br>50 (47.5-57.0) vs. 69 (63.5-71.8) | 4.017 | <0.001*** |
| Control <60s vs. AD<br>50 (47.5-57.0) vs. 74 (72.0-77.2) | 5.543 | <0.001*** |
| Control <60s vs. ALS<br>50 (47.5-57.0) vs. 70.5 (65.5-74.5) | 4.340 | <0.001*** |
| Control ≥60s vs. AD<br>69 (63.5-71.8) vs. 74 (72.0-77.2) | 1.813 | 0.210 |
| Control ≥60s vs. ALS<br>69 (63.5-71.8) vs. 70.5 (65.5-74.5) | 0.678 | 1.000 |
| AD vs. ALS<br>74 (72.0-77.2) vs. 70.5 (65.5-74.5) | 1.051 | 0.880 |

AD = Alzheimer's disease; ALS = Amyotrophic lateral sclerosis Median age at death is given in years, with 1<sup>st</sup>-3<sup>rd</sup> quartiles Overall group difference: $\chi^2=39.272$ , df=3, p<0.001; Kruskal-Wallis rank sum test Note: Dunn's post hoc test used for pairwise comparisons. p-values are adjusted \*p<0.001

### Tissue samples

All post-mortem tissue was collected via the Edinburgh Brain Bank (ethics approval from East of Scotland Research Ethics Service, (REC: 21/ES/0087, IRAS: 298880)) in line with the Human Tissue (Scotland) Act. Use of human tissue for post-mortem studies has been reviewed and approved by the Edinburgh Brain Bank ethics committee and the Academic and Clinical Central Office for Research and Development (ACCORD) medical research ethics committee (AMREC).

Formalin-fixed, paraffin-embedded (FFPE) tissue was cut on a Leica microtome into 4 μm thick serial sections, and were collected on Superfrost (ThermoFisher Scientific) microscope slides. Sections were baked overnight at 40°C before staining. Sections were dewaxed using successive xylene washes, followed by alcohol hydration and treatment with picric acid to minimise formalin pigment. Samples consisted of formalin-fixed-paraffin-embedded (FFPE) slides of three brain regions for each individual; amygdala, frontal cortex and hippocampus.

### Immunohistochemistry

To assess the presence of pathological TDP-43 in tissue samples, standard immunohistochemistry (IHC), and *in situ* hybridisation (ISH) was performed using; (i) TDP-43 RNA aptamer ^15,16^ to detect pathological TDP-43 in the context of toxic TDP-43 gain-of-function, and, (ii) *HDGFL2* cryptic exon BaseScope^TM^ *in situ* hybridisation (ISH) probe^1^ (Bio-Techne; 1236471-C1) to detect *HDGFL2* cryptic exons in the context of toxic TDP-43 loss-of-function Anti-ferritin antibody (Abcam; ab287968) was used to detect ferritin heavy chain/ferritin H subunit (a ferroxidase enzyme, encoded by the FHT1 gene which stores and releases iron) as a proxy for total iron in tissue.

#### TDP-43 RNA aptamer

Compared to currently available antibody approaches, the TDP-43 RNA aptamer offers increased sensitivity and specificity for pathological TDP-43 detection ^16,18^, comprehensive molecular characterisation^15^, comprehensive experimental validation (in silico, in vitro) ^15^, pathological benchmarking against widely used TDP-43 antibodies (e.g. pTDP-43, C-terminal) ^16^, and crucially, pathological benchmarking against TDP-43 loss-of-function cryptic exon (*HDGFL2* in this study, along with *Stathmin-2* ^16^) detection probes.

The TDP-43 RNA aptamer provides several advantages compared to antibody approaches, specifically including; (i) increased sensitivity facilitated by its small molecular size (approximately 2 kDa) allows for many more binding events to TDP-43 which adopts highly heterogenous populations of phase-separated assemblies with differing sizes and structures, (ii) increased sensitivity because it gets around buried isotopes as is often the case in aggregation-prone neurodegenerative disease-related proteins, (iii) increased specificity because it binds only pathological TDP-43, defined by a free RRM. Additionally, pathological TDP-43 binding occurs in a non-phosphorylation-dependent manner, indeed phosphorylated TDP-43 (pTDP-43), whilst a frequently encountered pathological post-translational modification (PTM) ^32^, is not the only pathological PTM and relying only on pTDP-43 can therefore miss the majority of pathological aggregation events, for example acetylated TDP-43 pathology ^33^, or TDP-43 aggregation in the absence of post-translational modification. Our recent work demonstrates that TDP-43 loss-of-function (cryptic exon detection) can be seen in the absence of pTDP-43 aggregates ^16^, however the burden of TDP-43 aggregates detected by the TDP-43 RNA aptamer corresponds with the burden of cryptic exons This is not solely down to PTM-agnostic recognition and can also be explained in part by the fact that aptamers are not subject to epitope masking. Epitope masking is one of the biggest limitations of antibody-based IHC, during tissue fixation (e.g., formalin fixation), cross-linking and structural changes can hide antibody binding sites. This often necessitates antigen retrieval (heat or enzymatic treatment) to unmask epitopes, which can be variable and sometimes damage the tissue or alter antigenicity. Aptamers are less affected due to their smaller size and aptamers can access recessed or partially masked binding sites that antibodies (being much larger) cannot ^34^. Aptamers also have different binding modes recognising 3D structural motifs rather than requiring a fully exposed linear epitope, making them less dependent on retrieval methods ^34^. Aptamers are also robust to fixation artifacts tending to maintain binding even when the target protein has undergone conformational changes from fixation, reducing the need for harsh retrieval steps ^34^.

The TDP-43 aptamer that we have employed in this work can discriminate between clinically relevant structural variants of TDP-43 ^35^. It can detect pathological TDP-43 in *SOD1* cases ^21^, validated by cryptic exon detection in the same cases ^16^, pathology that cannot be resolved using antibody-only approaches. This in addition to extensive experimental validation of TDP-43 binding sites, with experimental verified binding affinity (K_d_) quantification, and biophysical molecular dynamics characterisation for contact tightness, stability, and interface interactions (specifically, including the number of contacts, distance, and numbers of H-bonds to provide an estimate of the tightness of contacts, as well as average motion covariance information on the atomic fluctuations of the interface binding site) ^15^. Validation against an objectively appropriate negative control (reverse complementary RNA), and off-target amyloidogenic protein (e.g. Aβ42 and α-synuclein ^15^) has also been assessed. More generally the aptamer does not suffer from well appreciated disadvantages of antibodies including, batch effects, and production bottlenecks. Taken together, the combination of the TDP-43 aptamer with cryptic exon ISH offers an ideal dual-detection method to identify TDP-43 pathology with unprecedented accuracy.

The TDP-43 RNA aptamer is an *in silico* designed and experimentally validated RNA small molecule that binds pathological TDP-43, and whose application has been optimized and adapted for use for immunohistological staining of formalin fixed paraffin embedded (FFPE) human tissue. TDP-43 RNA aptamer tissue staining was carried out as described previously ^16,36^. Briefly, following deparaffinisation, rehydration, and 15-min picric acid treatment with subsequent water wash, antigen retrieval was carried out in citric acid buffer (pH 6) in a pressure cooker for 15-min, after which immunostaining was performed using the Novolink Polymer detection system (Leica Biosystems, Newcastle, UK). Following incubation with peroxidase block for 30-min, and a subsequent 5-min wash step with TBS, Avidin and biotin blocking steps were then carried out using a biotin blocking kit (ab64212), followed by a 5-min TBS wash step and a 5-min milli-Q water wash step. 156nM of TDP-43 RNA aptamer (TDP-43Apt CGG UGUUGCU with a 3’ Biotin-TEG modification, ATDBio, Southampton, UK) was prepared in Milli-Q water, applied to the tissue and incubated for 3-hr at 4 °C followed by incubation with 4% PFA overnight at 4 °C. A 5-min wash with Milli-Q water then preceded incubation with anti-biotin HRP antibody (ab6651) diluted 1 in 200 in milli-Q water for 30-min, followed by a 5-min wash step with Milli-Q water and subsequent incubation with DAB for 5 min. Slides were washed with water, counterstained with hematoxylin, “blued” with lithium carbonate, dried, and coverslipped with VectaMount.

#### HDGFL2 cryptic exon BaseScope^TM^ in situ hybridisation (ISH) probe

*HDGFL2* cryptic exon BaseScope^TM^ *in situ* hybridisation (ISH) probe^1^ staining was carried out as described previously, using BaseScope^TM^ RED Reagent Kits according to manufacturer’s protocols. In summary, following deparaffinisation, rehydration, and 15-min picric acid treatment with subsequent water wash, slides were allowed to air dry for 5-min. Endogenous peroxidases were blocked using a 3% hydrogen peroxide solution for 10-min at room temperature, followed by two water wash steps. Antigen retrieval was carried out in a steamer for 15 mins, using ACD RNAScope Target Retrieval Reagents. The slides were incubated in Protease III solution for 30 min at 40°C, and after two water wash steps, incubation in probe solution for 2-hr for hybridisation at 40°C in a HybEZ II Oven was carried out. Amp reagents 1–8 were applied according to the protocol. Slides were incubated in Fast Red for 10 min, counterstained with hematoxylin, “blued” with lithium carbonate, dried, and coverslipped with VectaMount.

#### Anti-ferritin antibody

Anti-ferritin antibody (Abcam; ab287968) immunostaining was carried out using the Novolink Polymer detection system (Leica Biosystems, Newcastle, UK). Following deparaffinisation, rehydration, and 15-min picric acid treatment with subsequent water wash, antigen retrieval was carried out in Tris-EDTA buffer (pH 9) in a pressure cooker for 10-min. Manufacturer’s immunostaining protocols were then carried out with overnight anti-ferritin antibody incubation at 4°C. Slides were washed with water, counterstained with hematoxylin, “blued” with lithium carbonate, dried, and coverslipped with VectaMount.

### Quantification, pathology grading and imaging

Absence and presence of TDP-43 pathology correspond with symptom absence and presence, albeit with exceptions in as many as a quarter of cases, so-called mismatch cases. However, absolute burden has never been shown to correlate with symptoms ^37^, for TDP-43. Therefore, common practice dictates a binary assessment of presence or absence of TDP-43 aggregates as assessed by a clinical pathologist as well as a quantitative digital pathology evaluation performed using QuPath superpixel analysis as performed previously ^38^. All assessors were blinded to all demographic and clinical information. In brief, mean DAB intensity for each case stained with TDP-43 RNA aptamer or ferritin antibody was digitally acquired by superpixel analysis using digital burden scoring performed using the freely available QuPath software implementing superpixel analysis using code published previously ^22,38^.

## Statistical methods

Statistics were carried out using R ^39^ (version 4-5.1).

The normality of continuous variables (e.g., age) was assessed using the Shapiro–Wilk test (*shapiro.test* function, *stats* ^39^ package). Homogeneity of variances was evaluated using Levene’s test (*leveneTest* function, *car* package).

For comparison of continuous variables across multiple groups, parametric One-Way ANOVA was used (*aov* function, *stats* ^39^ package) when normality and homogeneity of variances assumptions were met, followed by Tukey’s Honest Significant Difference (HSD) post hoc tests (*TukeyHSD* function, *stats* ^39^ package) for pairwise comparisons, controlling the family-wise error rate. When normality and homogeneity of variances assumptions were violated, non-parametric Kruskal–Wallis tests (*kruskal.test* function, *stats* ^39^ package) were applied, followed by Dunn’s post hoc tests (*dunn.test* function, *dunn.test* ^40^ package), with Bonferroni correction, which account for the global rank distribution and variance across all groups in pairwise comparisons, rather than performing isolated pairwise rank comparisons as with Wilcoxon rank-sum test.

Where rank-based Dunn’s tests were not suitable due to tied values or nearly identical rank distributions, to robustly assess pairwise differences between groups (<60s, *≥*60s, AD, ALS), we implemented a permutation-based testing framework using the *coin* ^41,42^ package in R - For each pair of groups, the observed test statistic was calculated from the data, and a null distribution was generated by randomly permuting the group labels 10,000 times and recalculating the test statistic for each permutation. The p-value for each comparison was obtained as the proportion of permutations in which the simulated statistic was at least as extreme as the observed value. In addition to permutation p-values, Cliff’s delta for each pairwise comparison was calculated (using R’s *effsize* ^43^ package) to quantify effect sizes, representing the probability that a randomly selected observation from one group exceeds a randomly selected observation from the other, providing an interpretable measure of magnitude and practical relevance alongside statistical significance.

Frequency-based tests of associations between categorical variables within and between groups were assessed using Pearson’s chi-square test (*chisq.test* function, *stats* ^39^ package), with Fisher’s exact test (*fisher.test* function, *stats* ^39^ package) applied when expected cell counts were small.

Linear regression models were fitted using l*m()* to examine the relationship between Ferritin intensity (predictor) and TDP-43 intensity (outcome).

Confidence intervals for TDP-43 brain pathology prevalence (reported here as percentages or proportions) were computed using the modified *Wald* method of Agresti and Coull^44^ (*binom.confint* function, binom ^45^ package). The Agresti-Caffo ^46^ method was used to compute confidence intervals for the difference between two proportions, and to test for a difference between them (implemented manually, as the Agresti-Caffo has no implementation in current R packages).

The *tidyverse* ^47^ package was used for data manipulation and visualisation; specifically, data summaries were computed using *dply r*^48^ summary functions (and base R ^39^), data were visualised and plotted using *ggplot2* ^49^, *ggpubr* ^50^, *ggtext* ^51^, *cowplot* ^52^, and *ComplexUpset* ^53^.

## Results

### Pathological TDP-43 gain- and loss-of-function occurs in all 3 brain regions in ageing and disease

#### TDP-43 histology in *≥60s* compared to <*60s*

Using TDP-43 RNA aptamer and an *HDGL2* cryptic exon BaseScope^TM^ *in situ* hybridisation (ISH) probe, we were able to identify affected and unaffected individuals with high sensitivity in our ageing cohorts.

Amongst affected individuals, pathological TDP-43 was detectable with RNA aptamer in amygdala, hippocampus and frontal cortex in both the <*60s* and *≥60s* cohorts (**Figure 1A**). This pathology represented true TDP-43 loss-of-function as indicated by the presence of *HDGFL2* cryptic exons where TDP-43 neuronal pathology was evident (**Figure 1B**).

#### TDP-43 histology in ALS and AD

ALS cases demonstrated widespread neuronal pathology across all regions. Contrastingly, but expectedly, AD cases exhibited both unaffected and affected brain regions across the cohort (**Figure 2**). Strikingly, relative TDP-43 burden (both intensity and distribution) did not differ between disease and control subgroups, with a greater burden of pathology detected compared to antibody approaches, corresponding well with the distribution of *HDGFL2* cryptic exons, as previously reported ^16^.

**Figure 2.**
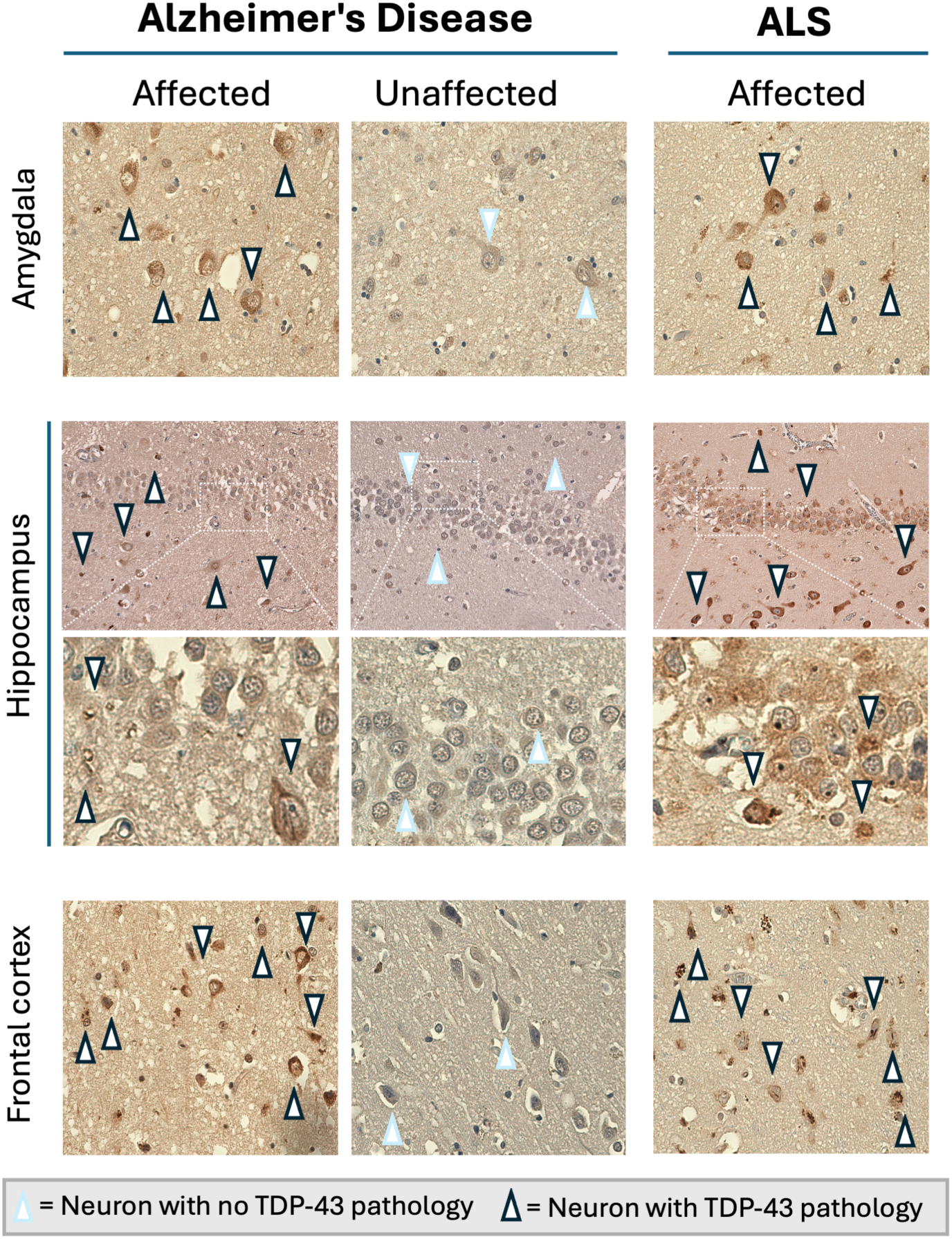
TDP-43 pathology occurs in all 3 brain regions in neurodegenerative disease. Representative images of the amygdala, hippocampus, and frontal cortex from unaffected (AD cases with no TDP-43 pathology) and affected (AD cases with TDP-43 pathology) AD and ALS cases (all had TDP-43 pathology) are shown. TDP-43 aptamer staining highlights neuronal pathology: neurons with no TDP-43 pathology are indicated by arrowheads with a light blue outline, whereas neurons exhibiting TDP-43 pathology are indicated by arrowheads with a black outline. Comparisons across brain regions and disease states reveal distinct patterns of TDP-43 pathology in AD and ALS, but with equivalent pathological burden in affected individuals.

#### TDP-43 RNA aptamer detection shows 1:1 correspondence with *HDGFL2* cryptic exon detection in a representative subset of ageing, AD and ALS cohorts

To investigate if TDP-43 loss-of-function (LOF) could be detected in amygdala, hippocampus and frontal cortex, and further demonstrate that TDP-43 RNA aptamer gain-of-function (GOF) detection corresponds with, and can be used as a proxy for TDP-43 LOF (see ^16^) we stained a representative subset tissue slides from each of our cohorts with *HDGFL2* BaseScope^TM^ ISH probes ^1^.

This representative subset consisted of 18 slides (12 positives and 6 negatives), the 12 positives included one slide for each of our 3 brain regions (i.e. amygdala, hippocampus and frontal cortex) across all 4 of our study cohorts (i.e. *<60s*, *≥60s*, AD, and ALS) and the negatives included one slide for each of the brain regions in the <60s and *≥*60s groups.

In all 12 instances, where TDP-43 RNA aptamer positive aggregates were detected, *HDGFL2* cryptic exon TDP-43 LOF was detected with BaseScope^TM^ ISH probes, with negative TDP-43 RNA aptamer staining corresponding to a lack of BaseScope^TM^ ISH staining for the 6 negative slides.

*HDGFL2* cryptic exons can be directly visualized using BaseScope single molecule ISH, showing a red dot wherever there is an mRNA containing the cryptic exon. Representative images show intracellular signal in neurons (**Figure 1B**, cell on right of image far right of panel) and glial cells (**Figure 1B**, cell on left of image far right of panel). Here we demonstrate TDP-43 LOF through the detection of *HDGFL2* cryptic exons in amygdala, hippocampus, and frontal cortex, confirming and extending previous TDP-43 LOF detection in hippocampus and motor cortex using a HDGFL2 cryptic peptide antibody ^13^.

## Hippocampal TDP-43 pathology is a specific signature of ageing

### TDP-43 pathology is a feature of normal ageing from the fourth decade, characterised by a 4.5-fold increase in hippocampus pathology in *≥60s* compared to <*60s*

#### TDP-43 brain pathology is a feature of normal ageing from the fourth decade

Amongst all non-neurodegenerative disease brains (i.e. brain bank controls) constituting the ageing cohorts, TDP-43 pathology was detected in 54.5% (18/33; CI 38.0-70.2%) of individuals (**Figure 3A**, **Table 3**).

**Figure 3.**
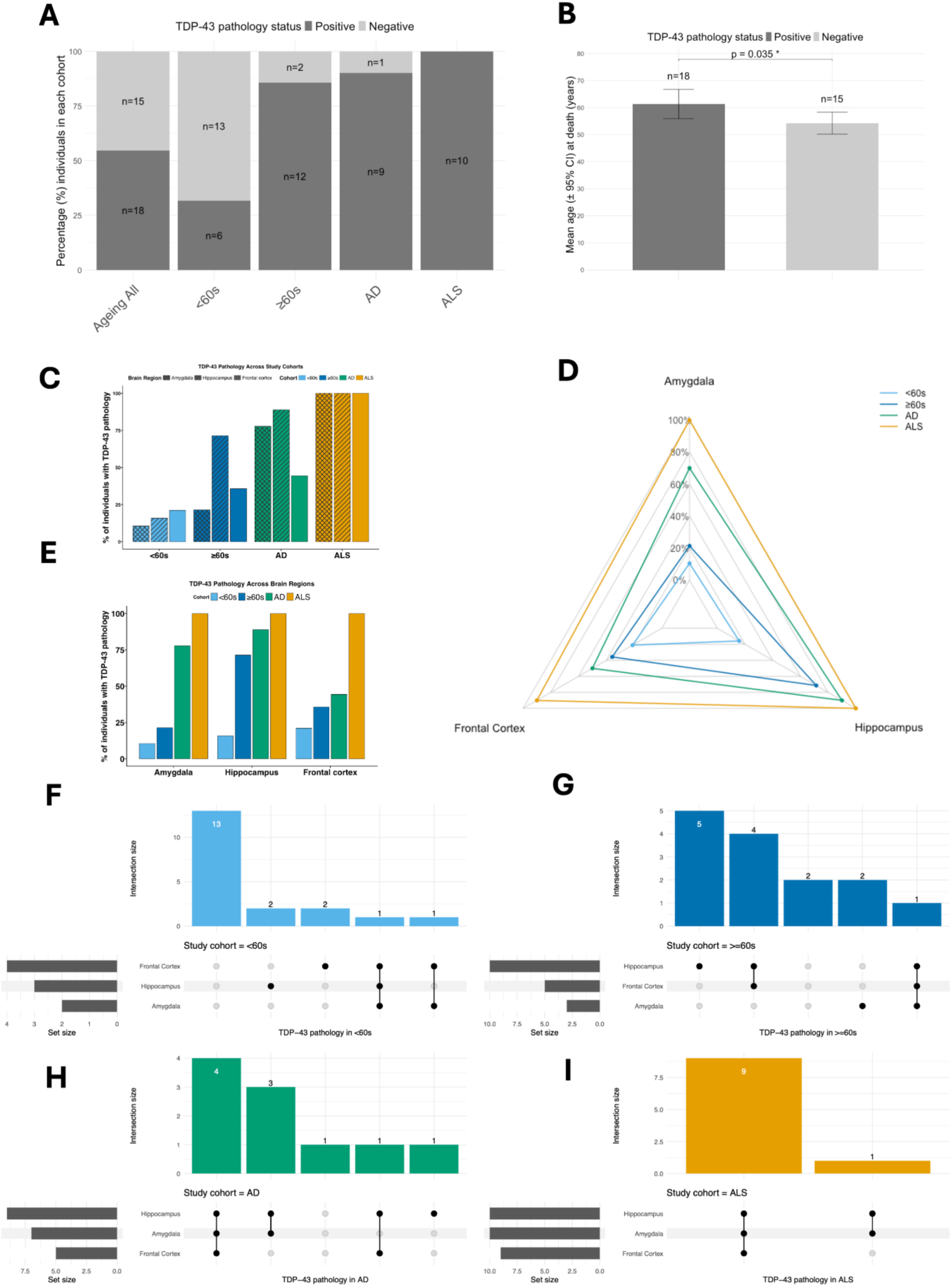
TDP-43 pathology is a feature of normal ageing, with brain region-specific signatures in AD and ALS. **A,** TDP-43 brain pathology prevalence amongst individuals in ageing (combined, <60s and ≥60s), AD and ALS cohorts, presented as percentages (%), and numbers of individuals with and without brain pathology. **B,** TDP-43 pathology prevalence is higher in ≥60s than <60s. Mean age of TDP-43 positive and negative individuals is presented with 95% confidence intervals, and p-value indicating significant difference by Welch’s t-test. **C,** Percentages of individuals with TDP-43 pathology, presented for each brain region by study cohort groups (<60s, ≥60s, AD, ALS). **D,** Radar plot illustrating percentages of individuals with TDP-43 from each cohort (<60s, ≥60s, AD, ALS) with plot tips for each brain region. **E,** Percentages of individuals with TDP-43 pathology, presented for study cohort (<60s, ≥60s, AD, ALS) by brain regions group. **F,** Upset plot illustrating incidence of brain region-specific, and brain region co-incident, TDP-43 pathology amongst individuals in the <60s cohort. **G,** Upset plot illustrating incidence of brain region-specific, and brain region co-incident, TDP-43 pathology amongst individuals in the ≥60s cohort. **H,** Upset plot illustrating incidence of brain region-specific, and brain region co-incident, TDP-43 pathology amongst individuals in the AD cohort. **I,** Upset plot illustrating incidence of brain region-specific, and brain region co-incident, TDP-43 pathology amongst individuals in the AD cohort. Study cohorts are colour-coded throughout as follows; <60s = light blue, ≥60s = dark blue, AD = green, ALS = orange.

**Table 3.**
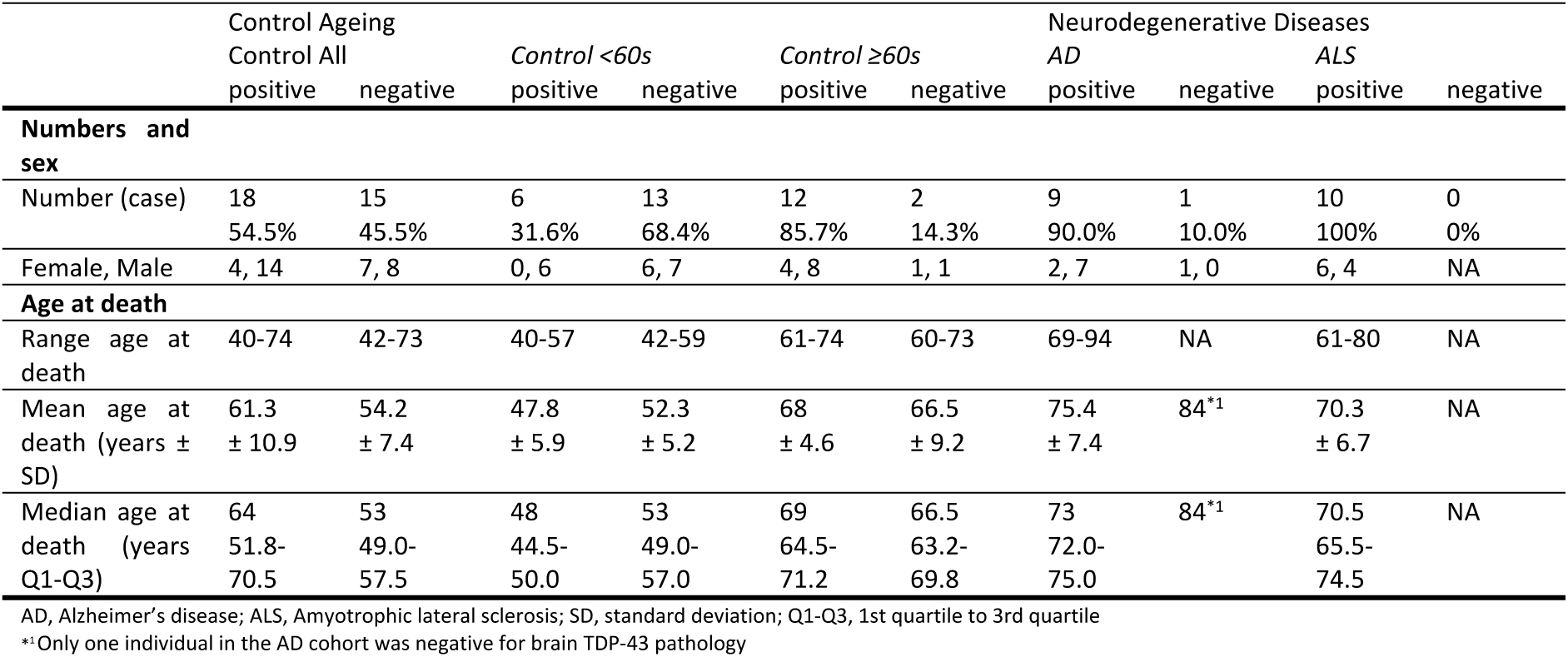
Presence of brain TDP-43 pathology with demographics across cohorts. TDP-43 pathology prevalence is indicated by the number of individuals positive or negative for brain TDP-43 in control ageing cohorts, including all controls and the <60s subgroup. For each subgroup of TDP-43 positive or negative, the number of cases, sex distribution, and age at death are presented.

Age at death for brain bank control individuals with TDP-43 pathology ranged from 40-74 years old, and was normally distributed (W=0.899, p=0.054; Shapiro test), with a mean of 61.3 (SD=10.9) years of age compared to that of 54.2 years (n=15, SD±7.4) for individuals without brain TDP-43 pathology (**Table 3**).

#### TDP-43 brain pathology incidence is increased with age

For control individuals, Shapiro-Wilk tests indicated no significant deviation from normality in age at death for individuals with TDP-43 aptamer pathology (W₁₈=0.899, p=0.054) or without (W₁₅=0.930, p=0.277). Levene’s test for homogeneity of variance, based on the median, indicated no evidence of unequal variances between the groups (F_1,31_=2.64, p=0.115), supporting the assumption of equality of variances for parametric testing. Nonetheless, giving the relatively small sample sizes of both groups, and a slight inequality in the number of individuals in both groups, we decided to eschew least conservative t-test approaches for differences in age at death between controls that were TDP-43 brain positive, and controls that were TDP-43 brain negative.

Using the more conservative parametric Welch’s t-test, we found a significant difference in mean age at death between ageing cohort individuals, with and without brain TDP-43 pathology (t=-2.211, df=29.818, p=0.035; **Figure 3B**). This finding was robust to parametric assumptions as indicated by non-parametric Wilcoxon rank sum test (equivalent to Mann–Whitney U test) (W=78.5, p=0.042)

#### TDP-43 brain pathology incidence is increased 2.7-fold between ≥60s and <60s cohorts

Amongst non-neurodegenerative disease brains examined, TDP-43 pathology was present in 31.6% (n=6/19; 95% CI 15.2-54.2) of individuals aged under 60 (the *<60s* cohort), and in 85.7% (n=12/14; 95% CI 58.8-97.2) of individuals greater than or equal to 60 years (the *≥60s* cohort) (**Figure 3A**, **Table 3**).

This difference in age represented a statistically significant 2.7-fold increase (z=3.3795, p<0.001; Agresti-Caffo z-test) in the incidence of brain TDP-43 pathology between individuals in the *≥60s* and *<60s* cohorts.

In the *<60s* cohort, the individual with brain TDP-43 pathology with earliest age at death was 40 years old, with TDP-43 pathology found in two further individuals in their forties (aged 44 and 46). Three individuals with autopsies in their 50s (aged 50, 50 and 57) showed pathological TDP-43. Notably, all six individuals under 60 (the *<60s* cohort) with TDP-43 pathology were male (**Table 3**).

Amongst the 12 individuals with TDP-43 brain pathology in the *≥60s* cohort, seven were in their 60s (2 females and 5 males; aged 61, 61, 63, 65, 67, 69, 69 at autopsy), and five were in their 70s (2 females and 3 males; aged 71, 71, 72, 73, 74 at autopsy).

#### Hippocampus TDP-43 pathology is a brain region-specific signature of ageing

TDP-43 pathology was present in all 3 brain regions (amygdala, frontal cortex and hippocampus) in both the *<60s* and *≥60s* cohorts, presenting as single-, double- or triple-brain region (co)pathologies in control ageing individuals.

In the younger <*60s* cohort, frontal cortex pathology was most common, detected in 21.1% (n=4/19; 95% CI 8.0-43.9; see **Figure 3C**, **Figure 3D**) of individuals. This was followed by hippocampus pathology present in 15.8% (n=3/19; 95% CI 4.7-38.4; **Figure 3C**, **Figure 3D**) of *<60s* brains, with amygdala pathology in 10.5% (n=2/19; 95% CI 1.7-32.6; **Figure 3C**, **Figure 3D**).

Amongst the six <*60s* individuals with TDP-43 pathology, 2 individuals had hippocampus only pathology, 2 had frontal cortex only pathology, 1 individual had amygdala and frontal cortex co-pathology, and the other individual had pathology in all 3 brain regions (**Figure 3F**).

In the older *≥60s* cohort, hippocampus pathology was present in 71.4% (n=10/14; 95% CI 45.0-88.7; **Figure 3C**, **Figure 3D**) of individuals, a statistically significant 4.5-fold increased incidence (z=3.445, p<0.001; Agresti-Caffo z-test) between *≥60s* and <*60s* cohorts (**Figure 3A3 3C**). Compared to <*60s*, whilst prevalence of amygdala pathology was increased 2-fold to 35.7% (n=5/14; 95% CI 16.2-61.4; **Figure 3C**, **Figure 3D**) in *≥60s*, this difference was not statistically significant (z=0.809, p=0.419; Agresti-Caffo z-test). Similarly, frontal cortex pathology prevalence was increased by 1.7-fold to 21.4% (n=3/14; 95% CI 6.8-48.3; **Figure 3C**, **Figure 3D**), this difference was also not statistically significant (z=0.897, p=0.370; Agresti-Caffo z-test). The dramatic (4.5-fold) increase in the incidence of TDP-43 pathology in hippocampus, compared to the other brain regions (amygdala and frontal cortex), comparing *≥60s* and <*60s* cohorts is further illustrated by radar plot (see **Figure 3E**).

Amongst the ten *≥60s* cohort individuals with TDP-43 pathology, 5 individuals had hippocampus only pathology, 2 individuals had amygdala only pathology, 4 individuals had hippocampus and frontal cortex co-pathology, and one individual had pathology in all 3 brain regions (**Figure 3G**).

#### Number of brain regions affected does not increase with age in control individuals

In controls, the increased incidence of TDP-43 pathology with age was not associated with an increased brain region involvement – i.e. in controls with TDP-43 pathology, we found only a weak, non-significant, negative association (ρ_33_=-0.256, p=0.305; Spearman’s rank correlation; **Suppl. Fig. 1B**), between age at death and the number of brain regions (1-3) with TDP-43 pathology.

Given that the number of brain regions affected with TDP-43 pathology does not increase with age in controls, we find no evidence to support spreading of TDP-43 brain pathology in ageing, amongst our cohort.

## AD is characterised by multi-region pathology, and increased amygdala pathology relative to age-matched individuals

### TDP-43 pathology is a common feature amongst individuals with AD

#### TDP-43 pathology is a very common feature amongst individuals with AD

For AD, 90% (9/10; 95% CI 57.4-100) of individuals had brain TDP-43 pathology (**Figure 3A**, **Table 3**). The median age at death for all AD cohort individuals was 74 (72.0-77.2) years old (**Table 1**), and 73 (72.0-75.0) for AD individuals with TDP-43 pathology (**Table 3**).

The single individual in the AD cohort without TDP-43 pathology was female and 84 years (oldest individual in the study) old at autopsy with a post mortem classification of “No significant abnormalities, Braak tangle stage 1, Moderate non-amyloid SVD”, raising the possibility of clinical misidentification in this instance.

#### AD is characterised by a >3-fold incidence of amygdala TDP-43 pathology relative to aged-matched individuals

Compared to individuals from the older *≥60s* cohort (i.e. an appropriate age-matched cohort; see **Table 2** for statistics), TDP-43 pathology incidence was increased in amygdala, hippocampus and frontal cortex amongst individuals in the AD cohort (see **Figure 3C**).

In AD for amygdala, a statistically significant 3.27-fold increase (z=2.3962, p=0.0166; Agresti-Caffo z-test) was observed relative to age-matched controls (**Figure 3C**).

These increases in incidence were not statistically significant for hippocampus (1.26-fold; z=0.9233, p=0.3564; Agresti-Caffo z-test) or frontal cortex (1.4-fold; z=0.6636, p=0.5069; Agresti-Caffo z-test), however (see **Figure 3C**).

The large (3.27-fold) increase in the incidence of TDP-43 pathology in amygdala in AD, compared to the other brain regions (hippocampus and frontal cortex), compared to younger <*60s*, and age-matched *≥60s*, cohorts is further illustrated by radar plot, here (**Figure 3D**).

TDP-43 co-pathologies across multiple brain regions is a characteristic hallmark of AD.

Among the 9 AD individuals with TDP-43 pathology, seven individuals had multi-region pathology – four individuals had pathology in all 3 brain regions, and 4 individuals had amygdala and hippocampus pathology (**Figure 3H**). One AD cohort individual had hippocampus only pathology. Frontal cortex TDP-43 pathology in AD was only observed in the context of triple brain-region co-pathologies (n=4; **Figure 3H**), perhaps suggesting that it may be a late brain region pathology in the context of widespread brain pathology.

## ALS is characterised by near universal triple-region TDP-43 pathology

### Triple-region TDP-43 pathology is widespread amongst individuals in the ALS cohort

#### TDP-43 pathology in all 3 brain regions is almost ubiquitous amongst individuals with ALS

All 10 individuals from the ALS cohort had TDP-43 brain pathology (**Figure 3A**). For 9 out of the 10 ALS cohort individuals with TDP-43 brain pathology, this pathology was always manifested in all three brain regions examined, i.e. amygdala, hippocampus, and frontal cortex (**Figure 3I**), consistent with our previous findings ^37^. The other individual had pathology in amygdala and hippocampus, but not in frontal cortex. The median age at death of individuals in the ALS cohort was 70.5 (65.5-74.5) years old (see **Table 1**).

### Absolute ferritin levels are brain region-specific in ageing and disease

Because brain iron stored in ferritin can be measured by MRI, we need to understand brain region differences in ageing and disease.

## Hippocampus ferritin is an AD-specific marker of neurodegenerative disease

### Amygdala ferritin levels are similar across ageing and disease

Amygdala median ferritin levels were significantly different between across all 4 cohorts (χ^2^=9.323, df=3, p-value=0.025; Kruskal-Wallis test). Whilst amygdala ferritin levels were higher in ≥60s than in <60s cohorts (**Figure 4A**), this difference was not statistically significant (δ= 0.353, p= 0.063; Permutation test).

**Figure 4.**
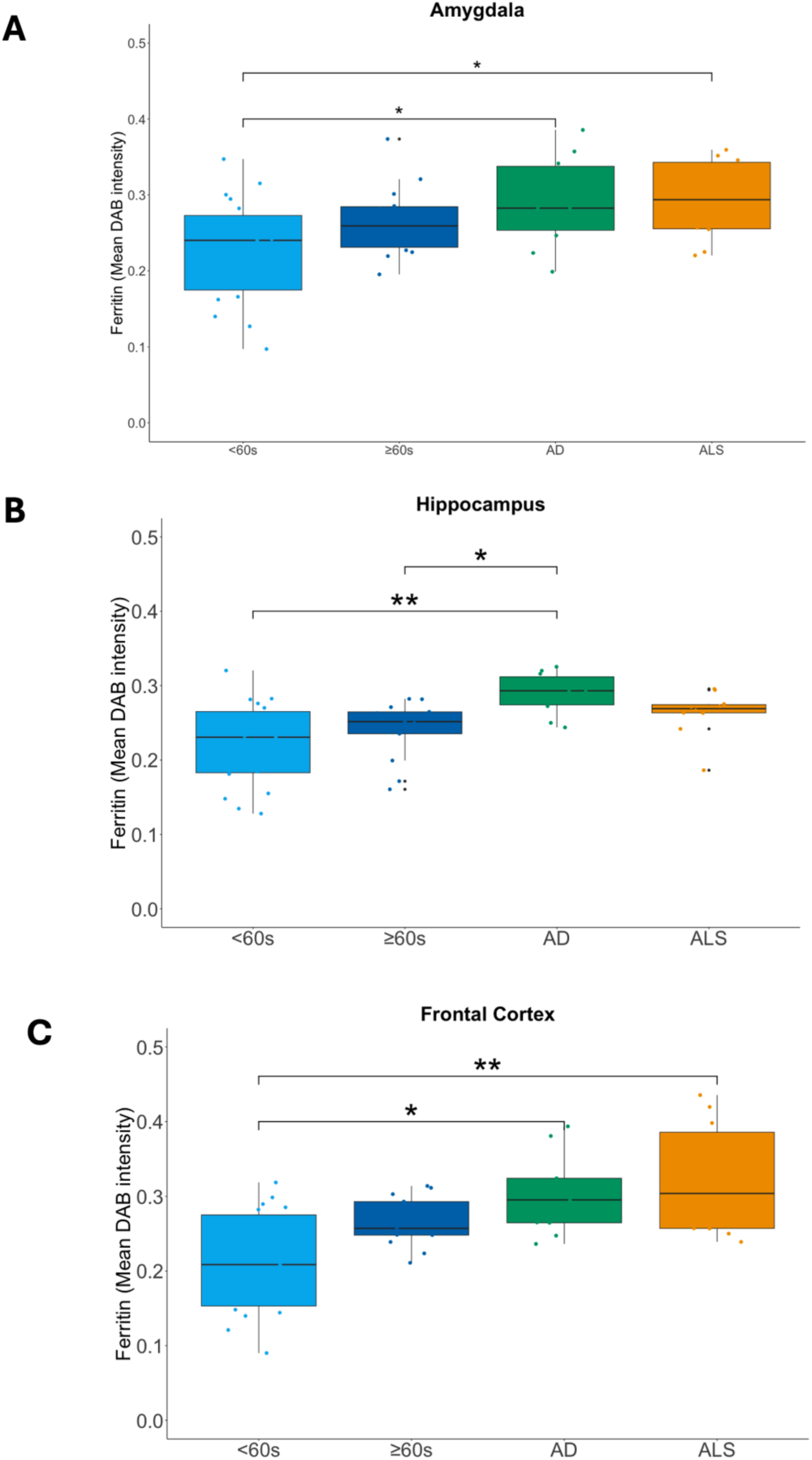
Absolute ferritin levels differ in ageing and disease in a brain region-specific manner, with high hippocampus ferritin a specific disease signature of AD. Boxplots indicating median ferritin levels from **A,** amygdala, **B,** hippocampus, and **C,** frontal cortex in <60s, ≥60s, AD, and ALS cohorts. Pairwise comparisons with significant differences in ferritin levels, as indicated by post hoc Dunn’s tests with Bonferroni correction, are illustrated with significance bars and indicative p-value asterisks (*p<0.05, **p<0.01).

Ferritin levels were also higher in disease (both for ALS and AD) than in ageing cohorts (**Figure 4A**), but no significant differences were observed in amygdala ferritin levels between ALS and age-matched controls (i.e. ≥60s) (δ= 0.300, p= 0.179; Permutation test), AD and age-matched controls (δ= 0.286, p= 0.235; Permutation test), or indeed between AD and ALS (δ<0.001, p=0.948; Permutation test). Amygdala median ferritin levels were significantly higher in AD compared to <60s (δ<0.547, p=0.013; Permutation test), and ALS compared to <60s (δ<0.537, p=0.011; Permutation test), but these cohorts were not age-matched (see **Table 1**, **Table 2)**.

### Higher hippocampus ferritin levels is a disease-specific signature of AD

Hippocampus median ferritin levels were significantly different between cohorts (χ^2^=14.308, df=3, p=0.003; Kruskal-Wallis test). For AD, significantly higher hippocampus ferritin levels were observed compared to age-matched (i.e. ≥60s) controls (z= 2.685, p= 0.022; Dunn’s test) (**Figure 4B**). Similarly, higher hippocampus ferritin levels were observed in AD compared to <60s (z= 3.550, p= 0.001; Dunn’s test) but as there was no significant difference in ferritin levels between <60s and ≥60s (z=0.716 =, p>0.999; Dunn’s test), the difference between AD and <60s could be attributable to age. Notably, hippocampus ferritin levels in ALS were lower but not significantly different to AD (z= 1.298, p= 0.582; Dunn’s test), or compared to age-matched ≥60s (z=1.303 =, p= 0.577; Dunn’s test) (**Figure 4B**).

### Higher frontal cortex ferritin levels in disease are likely a function of ageing

Frontal cortex median ferritin levels were significantly different between cohorts (χ^2^=14.254, df=3, p=0.003; Kruskal-Wallis test). Higher ferritin levels in disease were observed compared to both ageing cohorts (**Figure 4C**), but these differences were only significant between AD and <60s (z=2.861, p=0.013; Dunn’s test), and between ALS and <60s (z=3.273, p=0.003; Dunn’s test), but not for AD (z=1.1813, p=0.712; Dunn’s test) or ALS (z=1.506, p=0.396; Dunn’s test) compared to the age-matched ≥60s cohort.

Whilst, frontal cortex ferritin levels were higher in ≥60s than in <60s cohorts (**Figure 4C**), this difference was not statistically significant (z=1.793, p=0.219; Dunn’s test) indicating no evidence here for age-related increases in frontal cortex ferritin levels. Given our observation of highest ferritin levels in disease here, our findings tentatively suggest a possible age-disease interaction for frontal cortex susceptibility to increased ferritin levels but only for older people with disease (AD or ALS).

In addition to the analyses comparing ferritin levels for each of the three brain regions across all four cohorts (<60s, ≥60s, AD, ALS) together presented here, we analysed ageing- and disease-comparable groups separately (see **Supplementary information; Suppl Fig. 2**) but without significant differences in findings.

### Brain region ferritin is a viable biomarker for concurrent TDP-43 pathology

To investigate the potential of brain ferritin as a biomarker of concurrent TDP-43 pathology burden we examined the relationship between ferritin levels (predictor) and TDP-43 pathology burden (outcome) across brain regions in neurodegenerative disease and ageing.

## Brain region ferritin is a TDP-43 biomarker for distinguishing and stratifying disease

### Amygdala ferritin levels reflect TDP-43 pathology in AD and ALS

In the AD cohort, amygdala ferritin levels were strongly associated with TDP-43 pathology, with 66% of the variation in TDP-43 pathology burden explained by ferritin levels (R²=0.664, p<0.01; **Figure 5A**, **Table 4**), with a slope coefficient of 1.764 (**Figure 5B**, **Table 4**) indicating that for each unit increase in ferritin levels, TDP-43 pathology burdens increase on average by over 1.7 units.

**Figure 5.**
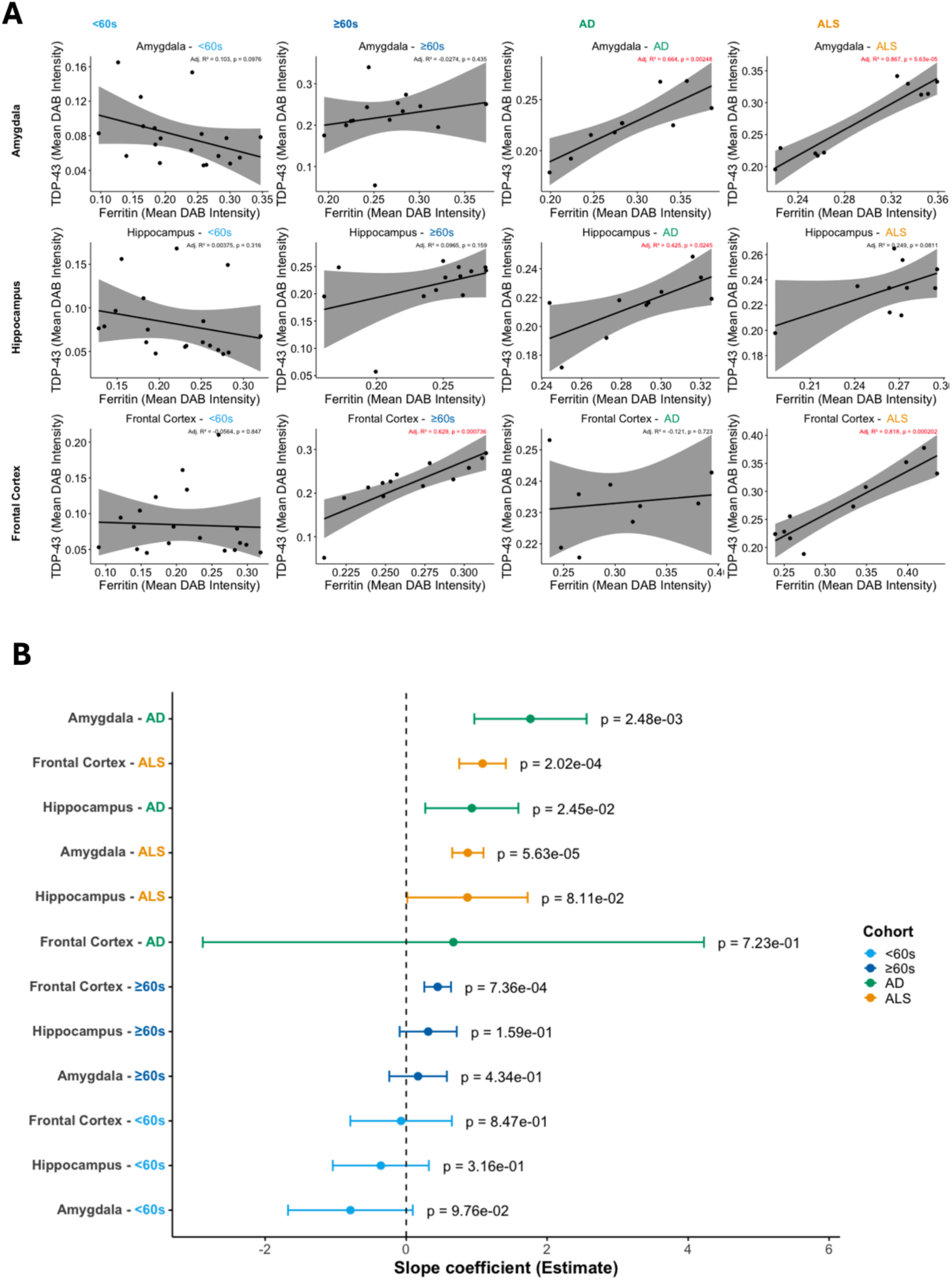
Regression analysis showing that ferritin levels predict TDP-43 pathology burden across specific brain regions in neurodegenerative disease and ageing. **A,** Scatterplots showing the relationship between ferritin levels and TDP-43 pathology burden across amygdala, hippocampus and frontal cortex brain regions in ageing (<60s and ≥60s), Alzheimer’s disease (AD), and amyotrophic lateral sclerosis (ALS). Each point represents an individual case, with colors indicating cohort. Regression lines with 95% confidence intervals depict the fitted relationship between ferritin (predictor) levels and TDP-43 burden (outcome). The adjusted R^2^ and p-value for the regression model are displayed. **B,** Forest plot of regression coefficient (with standard errors) and p-values from linear regression when ferritin levels are regressed against TDP-43 pathology burden for each cohort (<60s, ≥60s, AD, ALS) for amygdala, hippocampus and frontal cortex.

**Table 4.** Regression statistics assessing the contribution of ferritin variation to TDP-43 pathology burden. The table reports estimated regression coefficients, standard errors, confidence intervals, and significance levels for the association between ferritin levels and TDP-43 pathology across brain regions and clinical cohorts. Positive coefficients indicate higher TDP-43 burden with increasing ferritin levels, with significant results highlighting potential for brain region-specific ferritin levels as a biomarker for TDP-43 pathology burden in disease and ageing

| Cohort | Region | Linear regression model (Ferritin ~ TDP-43) statistics |  |  |  | Regression slope coefficients & p-values |  |  |
| --- | --- | --- | --- | --- | --- | --- | --- | --- |
|  |  | F | df | p-value (exact) | R<sup>2</sup> adjusted | Estimate | SE | p-value (significant) |
| <60s | Amygdala | 3.07 | 1,17 | 0.0976 | 0.103 | -0.791 | 0.45102 | 0.098 |
| ≥60s | Amygdala | 0.65 | 1,12 | 0.4345 | -0.027 | 0.168 | 0.2083 | 0.435 |
| AD | Amygdala | 18.82 | 1,8 | 0.00248 | 0.664 | 1.764 | 0.4066 | < 0.01** |
| ALS | Amygdala | 59.62 | 1,8 | 0.00006 | 0.867 | 0.875 | 0.11331 | < 0.001*** |
| <60s | Hippocampus | 1.07 | 1,17 | 0.316 | 0.004 | -0.359 | 0.34739 | 0.316 |
| ≥60s | Hippocampus | 2.28 | 1,11 | 0.15909 | 0.097 | 0.312 | 0.20688 | 0.159 |
| AD | Hippocampus | 7.64 | 1,8 | 0.0245 | 0.425 | 0.932 | 0.33724 | < 0.05* |
| ALS | Hippocampus | 3.98 | 1,8 | 0.0811 | 0.249 | 0.870 | 0.43587 | 0.081 |
| <60s | Frontal cortex | 0.04 | 1,17 | 0.847 | -0.056 | -0.072 | 0.36726 | 0.847 |
| ≥60s | Frontal cortex | 21.38 | 1,11 | 0.00073 | 0.629 | 0.447 | 0.09657 | < 0.001*** |
| AD | Frontal cortex | 0.14 | 1,7 | 0.723 | -0.121 | 0.670 | 1.8133 | 0.723 |
| ALS | Frontal cortex | 41.39 | 1,8 | 0.00020 | 0.818 | 1.084 | 0.16853 | < 0.001*** |
| Scatterplots with R<sup>2</sup> and p-values are presented in Figure 5A. |  |  |  |  |  |  |  |  |
| Forest plots of regression slope estimates with SE are presented in Figure 5B |  |  |  |  |  |  |  |  |

In the ALS cohort, this association was even stronger with 87% of the variation in TDP-43 pathology burden explained by ferritin levels (R²=0.867, p < 0.001; **Figure 5A**, **Table 4**), with a positive coefficient (β=0.875; **Figure 5B**, **Table 4**), demonstrating that amygdala ferritin levels explain a large proportion of TDP-43 pathology in ALS. Interestingly, for amygdala in the ALS cohort, two distinct populations of individuals seemed to present; one population with higher ferritin levels and corresponding high TDP-43 pathology, and a second with lower ferritin levels and corresponding lower TDP-43 pathology (see **Figure 5A**).

No statistically significant relationships were uncovered for either ageing cohort, with two important implications; that the linear, predictive relationship between ferritin and TDP-43 pathology in amygdala is disease-specific signature, not observed in younger control individuals or with ageing.

### Hippocampus ferritin is a viable biomarker for TDP-43 pathology in AD only

For hippocampus, a significant association and positive predictive relationship between ferritin levels and TDP-43 pathology burden was revealed for AD (R²=0.43, p<0.05; **Figure 5A**, **Table 4)** only (β=0.932; **Figure 5B**, **Table 4**).

### Frontal cortex ferritin reflects TDP-43 levels in age- and disease-specific manner

For frontal cortex, an age-dependent association between ferritin levels and TDP-43 pathology was revealed, with a significant association in the older ≥60s cohort (R²=0.63, p<0.0021; **Figure 5A**, **Table 4**) only (β=0.932; **Figure 5B**, **Table 4**), but not in <60s.

Indeed, for ALS, this frontal cortex association of ferritin levels and TDP-43 pathology burden was stronger, with 81% of the variation in TDP-43 pathology burden explained by ferritin levels (R²=0.818, p < 0.001; **Figure 5A**, **Table 4**), with a positive coefficient (β=1.084; **Figure 5B**, **Table 4**) indicating a near 1:1 linear relationship between changes in ferritin and TDP-43 pathology.

## Discussion

Our study provides the first estimate of the extent of TDP-43 brain pathology in ageing, ALS, and AD using an RNA aptamer with greater sensitivity compared to previous antibody-only approaches, revealing a hitherto underestimated extent of brain TDP-43 in this disease. Our findings demonstrate that TDP-43 pathology is not confined to specific neurodegenerative diseases but is also detectable across the normal ageing spectrum. We observed pathology in individuals as young as 40 years, with prevalence increasing steadily with advancing age. We demonstrate that in younger (age <60 years) but not older (age ≥60) individuals, that TDP-43 pathology corresponds to a lower age at death, implying divergent toxic mechanisms across life course. This supports the notion that TDP-43 is an age-associated pathology, extending earlier work that largely focused on older cohorts. Notably, our estimates suggest a somewhat higher prevalence than previous reports of phosphorylated TDP-43 in ageing populations ^2^, reflecting both methodological differences and our inclusion of younger individuals who were not represented in earlier studies. Regionally, TDP-43 pathology in ageing was most frequent in the hippocampus (80% of individuals aged ≥60 years) and the frontal cortex (40%), consistent with prior evidence that limbic and cortical regions are especially vulnerable. Importantly, none of the control individuals met criteria for limbic-predominant age-related TDP-43 encephalopathy (LATE), as no cognitive impairment was documented during life. This distinction underscores the difference between incidental TDP-43 pathology in ageing and clinically manifest disease.

The presence of TDP-43 brain pathology without a neurological and/or neuropsychiatric diagnosis, seemingly as a consequence of normal ageing suggests a number of non-exclusive hypotheses regarding the relationship between TDP-43 brain pathology and disease, i.e. that; (i) resilience factors such as high protein misfolding tolerance levels (e.g. chaperone-mediated) play a part in determining whether individuals develop symptoms and disease, (ii) TDP-43 brain pathology is in itself not-sufficient to cause disease but in combination with other neurotoxic factors (e.g. additional concomitant proteinopathies or heavy metal accumulation in the accumulation of damage following concomitant oxidative stress or mitochondrial damage) leads to disease, (iii) TDP-43 is not causative of disease but in fact a biomarker of disease in an underappreciated strong association with another proteinopathy (e.g. FUS, Annexin A11) whose pathological effect in ALS is also underappreciated or historically understudied.

In AD cases that we profiled, we observed the same age-related pattern of hippocampal and frontal cortical involvement as in controls ≥60 years, but with an additional, disease-specific signature of amygdala pathology. Classically AD is associated with hippocampus pathology driven by Aβ and/or tau burden, our findings here suggest brain region-specific susceptibility to TDP-43 pathology also, with increased amygdala involvement in AD compared to age-matched controls, which could be driving mood disorders frequently observed with AD. Strikingly, 80% (8/10) of AD cases exhibited hippocampal TDP-43 pathology, despite none being diagnosed with hippocampal sclerosis at post-mortem. This finding highlights the presence of TDP-43 pathology independent of overt hippocampal sclerosis and raises the possibility that hippocampal TDP-43 may contribute to cognitive decline in AD through mechanisms distinct from sclerosis. In ALS, all 10 cases showed TDP-43 pathology in all three regions examined (the hippocampus, frontal cortex, and amygdala) demonstrating that TDP-43 involvement extends well beyond the motor cortex. This widespread distribution underscores the multisystem nature of ALS pathology and may help explain the cognitive and behavioral features increasingly recognized in this disease.

Our recent work has highlighted the role of markers of redox dysfunction in ALS demonstrating that heavy metal accumulation in the form of iron accumulated ferritin in specific brain regions could be a viable biomarker for concurrent TDP-43 pathology ^22,54^. Our extended analysis in this cohort indicates that ferritin levels in distinct brain regions represent viable biomarkers for TDP-43 pathology, with potential utility in both disease detection and stratification. Specifically, brain region-specific ferritin is a viable biomarker for concurrent TDP-43 pathology and can distinguish and stratify disease. Amygdala ferritin levels reflect TDP-43 pathology in both AD and ALS, while hippocampal ferritin appears to be a specific marker for TDP-43 pathology in AD. This agrees with previous imaging studies which reveal increased ferritin iron in conjunction with R2 signal in the hippocampus of AD patients compared to non-neurological controls, highlighting the importance of this signature in future differential diagnostic imaging development ^55^. In the frontal cortex, ferritin levels correspond to TDP-43 burden in an age- and disease-specific manner, suggesting regional variability in its diagnostic relevance. Indeed, we previously demonstrated that in ALS, amygdala ferritin and TDP-43 pathology was higher in those with behavioural deficit ^22^, this may be the case with the two groups formed in the current study. Collectively, these results support the use of region-specific measurements of ferritin accumulated iron, for example using iron-sensitive sequences on MRI, especially susceptibility weighted imaging (SWI) and T2*-weighted imaging (T2*WI), as a tool for detecting, differentiating, and characterizing TDP-43-related pathology across neurodegenerative conditions.

Given our findings, and in cognizance of recent developments and investigations using cryptic exons and peptides, we suggest that the adaptation of (i) TDP-43^APT^ for assessing TDP-43 toxic gain-of-function, and/or (ii) cryptic exon/peptide analysis for assessing TDP-43 toxic loss-of-function, holds significant potential for uncovering previously undiscovered or underappreciated age-related TDP-43 pathologies, irrespective of the presence/absence of symptoms. This could have important implications for understanding the etiology and ontogeny of neurodegenerative diseases, as well as for diagnostics and therapeutics.

Together, these results refine our understanding of TDP-43 pathology across ageing and disease. They suggest that while age-associated pathology is common and regionally selective, disease-specific signatures such as amygdala involvement in AD and widespread distribution in ALS distinguish pathological ageing from neurodegenerative processes. These findings emphasize the need for more sensitive detection methods and broader regional assessments to capture the true burden of TDP-43 pathology and to clarify its clinical significance.

## Contributors

*JMG* conceptualized, secured primary funding for, and led the study with key input from *FMW*, *HS*, *PCW*, and *JPL*. *JMG* and *HS* prepared the tissue request and coordinated sample logistics. Immunohistochemistry was performed by *HS*, *OST*, *FLR*, *FMW*, and *JMG*, with support from *PCW*, *JPL*, *IRS*, and *KEI*. *HS* and *JMG* undertook digital image acquisition and curation. Data curation was performed by *HS*, *FMW*, and *JMG*. Data analysis was performed by *FMW*, *HS* and *JMG*. *FMW* drafted the initial manuscript; subsequent drafts were prepared by all authors and finalized by *FMW*, *HS*, and *JMG*.

## Declaration of interests

The authors declare no competing interests.

## Data sharing

All data are available in the manuscript, tables and supplementary data.

## Funding

The research for this manuscript has been supported by: (i) a Target ALS Early-Stage Clinician Scientist award to *JMG* (FS-2023-ESC-S2), (ii) an NHS Grampian grant to *FMW* (GCA25107); (iii) an NIH grant to *JMG* and employing *HS* and *FR* (R01NS127186); a (iv) MND Scotland/CSO grant to *KR* (CAF/24/13 – 2024/MNDS/6500/780ROB); (v) a Motor Neuron Disease Association grant to *JMG* and employing *KH* (Gregory/APR24/2374-791); (vi) a LifeArc MND Primer Fund to *JMG* (A20170) (vii) an NIH grant RF1NS095969 to *PCW* (viii) and a Food & Drug Administration grant 1U01FD008129 to *PCW*. Funders had no role in study design, data collection, data analyses, interpretation, or writing the manuscript.

## Acknowledgments

The authors would like to thank the University of Aberdeen Microscopy and Histology Core Facility in the Institute of Medical Sciences, the Edinburgh Brain Bank and the patients and families of the donors without whom this study would not be possible.

## Supplementary Information

**Supplementary Figure 1.**
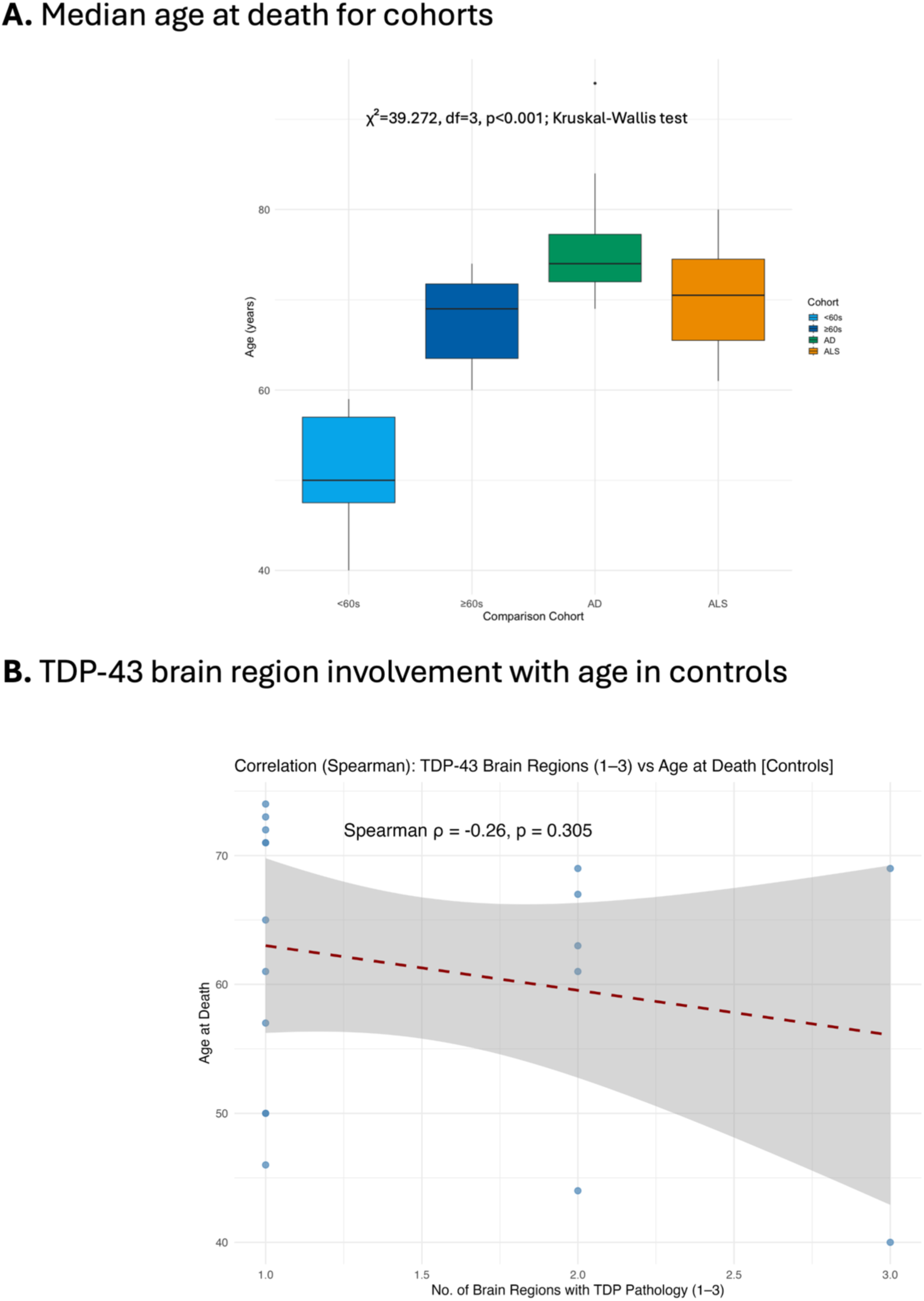
**A,** Median age at death boxplots for each of the four study cohorts (<60s, ≥60s, ALS and AD) (see also table 2). **B,** TDP-43 brain region involvement with age in ageing controls.

## Additional analysis

### Ferritin level differences are brain region-specific in ageing and disease

Because brain ferritin accumulated iron can be measured by MRI, we need to understand brain region differences in ageing and disease. In addition to the analyses comparing ferritin levels across all four cohorts (<60s, ≥60s, AD, ALS) together, we analysed ageing- and disease-comparable groups separately here, but without significant differences in results.

To do these analyses we tested brain-region specific ageing-related differences in ferritin levels between <60s and ≥60s (“Ageing Comparison), and disease-related differences in ferritin levels between AD, ALS and the age-matched ≥60s cohort (Disease Comparison).

No evidence for non-normality in ferritin level data was uncovered for <60s, ≥60s, AD or ALS cohort by Shapiro-Wilk tests, however inequality of variances amongst groups in the “Ageing comparison” between <60s and ≥60s (F=8.547, df=1,30, p=0.007; Levene’s test), and in the “Disease Comparison” between AD, ALS and ≥60s (F=5.1855, df=2,29, p=0.012; Levene’s test), informed our non-parametric approach here.

### Hippocampus ferritin is an AD-specific marker of disease

#### Amygdala ferritin levels similar across ageing and disease

Amygdala median ferritin levels were higher in ≥60s than in <60s but this difference was not significant (W=86, p=0.091; Wilcoxon rank sum test) (**Suppl. Fig. 2**). No significant difference in amygdala ferritin levels between AD, ALS or age-matched ≥60s (χ^2^= 2.059, df=2, p= 0.357; Kruskal-Wallis rank sum test) was revealed (**Suppl. Fig. 2).**

#### Higher hippocampus ferritin levels is a disease-specific signature of AD

Hippocampus median ferritin levels were higher in ≥60s than in <60s but this difference was not significant (W=151, p=0.305; Wilcoxon rank sum test) (**Suppl. Fig. 2**). However, hippocampus ferritin levels were significantly different in the disease comparison between AD, ALS, and age-matched ≥60s (χ^2^= 9.502, df=2, p= 0.009; Kruskal-Wallis rank sum test) (**Suppl. Fig. 2**). Here, for AD significantly higher hippocampus ferritin levels were increased compared to age-matched (i.e. ≥60s) controls (z= 3.08, p= 0.006; Dunn’s test).

Hippocampus ferritin levels in ALS were lower but not significantly different to AD (z= 1.54, p= 0.372; Dunn’s test), or significantly different compared to age-matched ≥60s (z=1.45 =, p= 0.455; Dunn’s test) (**Suppl. Fig. 2**).

#### Higher frontal cortex ferritin levels in disease are likely a function of ageing

Frontal cortex median ferritin levels were higher in ≥60s than in <60s (W=180, p=0.030; Wilcoxon rank sum test) (**Suppl. Fig. 2**), however but not significantly different in the disease comparison between AD, ALS or age-matched ≥60s cohorts (χ^2^=4.094, df=2, p= 0.129; Kruskal-Wallis rank sum test).

**Supplementary Figure 2:**
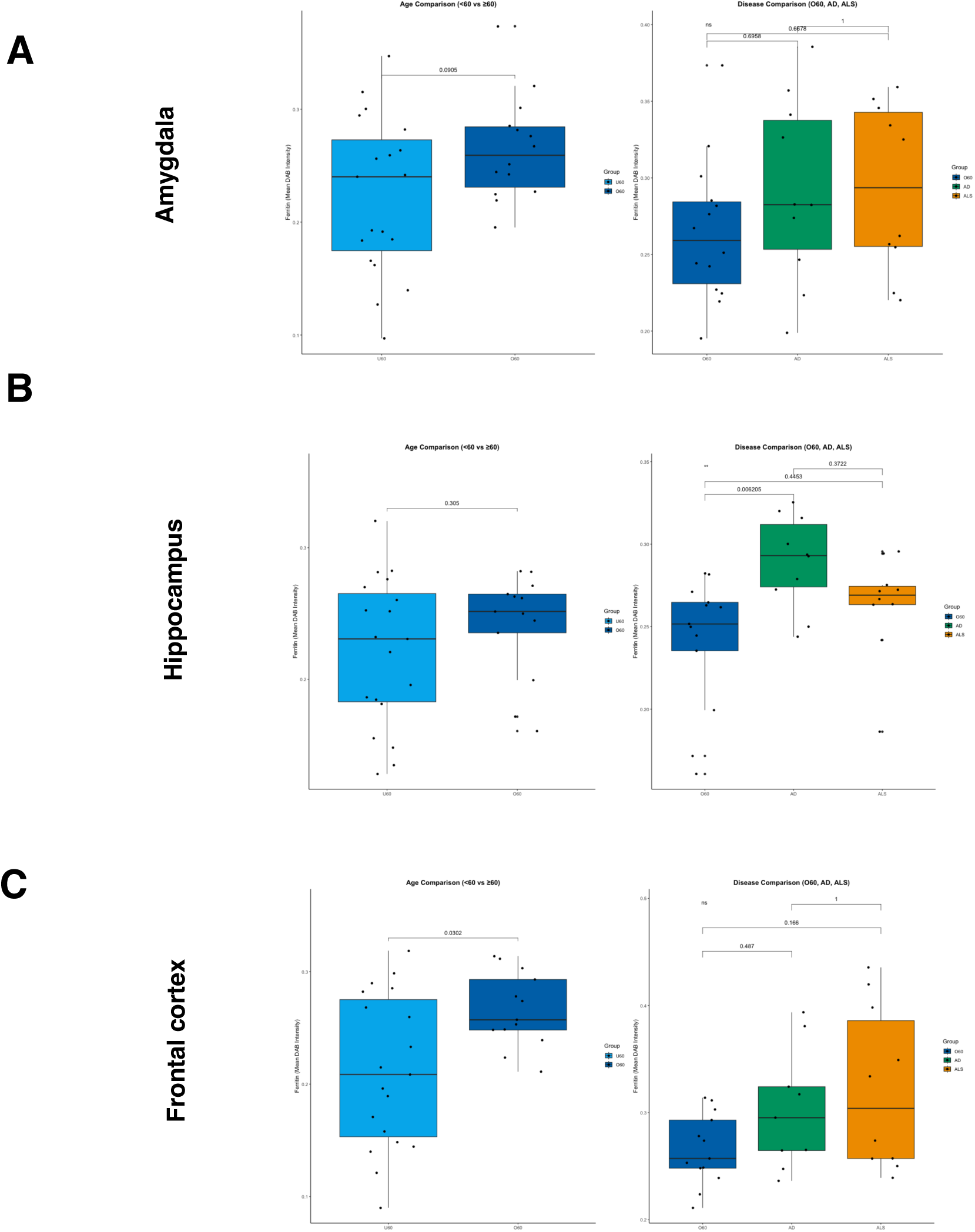
Increased hippocampus ferritin levels are a disease-specific signature of AD. For amygdala (A), hippocampus (B) and frontal cortex (C), median ferritin levels were statistically compared between <60s and ≥60s (“Ageing Comparison), and AD, ALS and the age-matched ≥60s cohort (Disease Comparison) using Wilcoxon rank sum tests and Kruskal-Wallis rank sum tests, respectively with p-values for group comparison presented with significance bars.

## Notes

### Competing Interest Statement

The authors have declared no competing interest.

